# Maternal cannabinoid exposure during lactation alters the developmental trajectory of prefrontal cortex GABA-currents in offspring

**DOI:** 10.1101/336735

**Authors:** Andrew F. Scheyer, Jim Wager-Miller, Anne-Laure Pelissier-Alicot, Michelle N. Murphy, Ken Mackie, Olivier J.J. Manzoni

## Abstract

Cannabis is the most widely used illicit drug in the world, and its usage is increasing with its widespread legalization. Use of the drug by mothers during lactation may transfer active cannabinoids to the developing offspring, altering postnatal neurodevelopment during this critical period. During early life, GABA undergoes a functional switch from an excitatory to an inhibitory neurotransmitter due to reciprocal changes in expression of the K^+^/Cl^-^ co-transporters KCC2 and NKCC1. Here, we characterize the functional GABA switch in the prefrontal cortex of both male and female rats. We show that treating rat dams with Δ9-THC or a synthetic cannabinoid during early lactation (PND01-10) retards KCC2 expression and delays the GABA switch in pups of both sexes via a CB1R-dependent mechanism. Our results indicate that the developmental trajectory of GABA in PFC neurons is significantly altered by perinatal exposure to cannabinoids through lactation during the early perinatal period.

## Introduction

Cannabis is the most widely used illicit drug in the world, and its usage in Western nations, including the United States continues to increase (United Nations Office on Drugs and Crime 2017). The best-characterized actions of cannabis are primarily attributed to Δ9-tetrahydrocannabinol (THC) and its effects on the endogenous cannabinoid receptors (CBR1 and CBR2), which together with naturally occurring endocannabinoids (eCBs) and their synthesizing and degrading enzymes, comprise the endogenous cannabinoid system (ECS; Bari et al. 2006). Consumption of cannabis during pregnancy is reported as between 1-6% (Metz and Stickrath 2015; Fergusson et al. 2002) and will likely rise as an increasing number of states and countries decriminalize or legalize its use. Public perception continues to categorize cannabis usage during pregnancy as of little-to-no consequence (Jarlenski et al. 2016) however, little is known about the transfer of THC and other active constituents of cannabis through breast milk, or the impact of the resultant neonatal exposure.

The role of the ECS during early development has been well-established in both animals (Trezza et al. 2012; Fride 2004) and humans (Fride et al. 2009; Mato, Del Olmo, and Pazos 2003; Mulder et al. 2008). Importantly, it is known that the primary active constituent of cannabis, THC, readily crosses the placental barrier and is found in breast milk (Perez-Reyes and Wall 1982; Marchei et al. 2011). Previous research indicates that exposure to THC has adverse consequences on fetal and perinatal neurodevelopment (Astley and Little 1990; English et al. 1997; Gunn et al. 2016; Richardson, Hester, and McLemore 2016) which may alter the course of development throughout the life cycle (Trezza et al. 2012; Campolongo et al. 2011). Furthermore, the ECS serves a crucial role in the development of the prefrontal cortex (PFC; Dow-Edwards and Silva 2017), a functional cognitive hub whose perturbation during early development has been linked to a variety of profound deficits in maturity (Goldstein and Volkow 2011; Schubert, Martens, and Kolk 2015; Scheyer, Martin, and Manzoni 2017).

The PFC is the most highly evolved mammalian brain region (Araque et al. 2017; Scheyer, Martin, and Manzoni 2017), functionally participating in a variety of behaviors from working memory and action planning to cognitive flexibility and emotionally-guided behaviors (Euston, Gruber, and McNaughton 2012; Goldman-Rakic 1991). The ECS functions as a modulatory neurotransmitter system in the PFC, highly concentrated at interneuron synapses (Berghuis et al. 2007; Mulder et al. 2008) and more prevalent in deep than superficial layers (Fortin and Levine 2007; Heng et al. 2011). Importantly, eCBs serve a critical function in the developmental trajectory of GABAergic interneurons (Berghuis et al. 2005). Consequently, perturbation of the ECS during neonatal development has lasting effects on GABAergic transmission (Garcia-Gil et al. 1999).

GABAergic synapses undergo significant development during early life. While GABA serves as the primary inhibitory neurotransmitter in the adult brain, its role is rather as an excitatory neurotransmitter in the immature brain due to a comparatively high intracellular Cl^-^ concentration caused by low levels of the KCC2 chloride transporter (Kaila et al. 2014). During development, GABA’s role undergoes a critical shift from excitatory to inhibitory, which is mediated by a large increase of KCC2 expression and a more modest increase in NKCC1, with a subsequent decrease in intracellular Cl^-^ (Kaila et al. 2014; Rivera et al. 1999; Ben-Ari 2002; Ben-Ari et al. 2007). The timing of this transition is a crucial point in the neurodevelopmental trajectory and aberrations during this critical period are linked with a number of neurological disorders including autism, Downs syndrome, Fragile X and schizophrenia (Ben-Ari 2008; Garber 2007; Zoghbi 2003). The exact timing of this transition differs between brain regions, beginning early as E15.5 in the hippocampus and as late as P15 in regions of the neocortex (Watanabe and Fukuda 2015). In the PFC, development of GABA synapses is maximal between P10 and P15 (Virtanen et al. 2018), though the functional valence of these sites has yet to be investigated.

There have been few investigations into the consequences on GABAergic function of perturbing the ECS during the critical postnatal neurodevelopmental period, though what data do exist suggest that the impact is significant and lasting (Schneider 2009). Indeed, while it is known that cannabis exposure during PFC development has significant, lasting effects (Realini, Rubino, and Parolaro 2009), the mechanistic underpinnings of these effects remain largely unexplored. Here, we investigated the impact of cannabinoids via maternal exposure during early postnatal life in order to assess the potential risks associated with cannabis use during this period. Importantly, the first two weeks of postnatal development in the rat are analogous to third-trimester development in the human (Khazipov and Luhmann 2006; Semple et al. 2013). Therefore, by investigating the impact of exogenous cannabinoid exposure during this period, we address both their impact on late in utero development and the impact of cannabinoids transferred through breast milk on the neonatal brain. Additionally, we established a time-course for the developmental trajectory of GABA’s transition from an excitatory to inhibitory neurotransmitter in the PFC. Together, these findings offer valuable insight into the potential risks of cannabis use during both late in utero development as well as during lactation on the developing brain.

## Results

### GABA transitions from an excitatory to inhibitory neurotransmitter in the PFC between P10 and P15

First, in order to establish the timing of GABA’s transition from an excitatory to inhibitory neurotransmitter in the rat mPFC, we used cell-attached recordings in mPFC slices of both male and female rats containing layer 5 pyramidal neurons to observe spontaneous cell spiking activity before and after the application of either the GABAR antagonist picrotoxin (PTX) or the GABAR positive allosteric modulator isoguavacine (ISO) as previously described (Ben-Ari et al. 2007; Lagostena et al. 2010; Figure 1). At P09-P10, application of PTX induced a significant decrease in spike frequency as compared to baseline (∼30% reduction; P<0.05 as compared to baseline). Conversely, application of ISO significantly increased (∼30% increase; P<0.05 as compared to baseline) spike activity as compared to baseline. These results are compatible with the idea that GABA serves as an excitatory neurotransmitter at this early time point. Conversely, cells recorded between P15-P16 exhibited increased spiking activity following PTX application (∼53% increase; P<0.05 as compared to baseline) while application of ISO significantly attenuated (∼39% reduction; P<0.05 as compared to baseline) spike frequency. Thus, at this later time point GABA functions as an inhibitory neurotransmitter. Together, these results indicate that in layer 5 pyramidal cells of the PFC, GABA undergoes a functional “switch” from an excitatory to an inhibitory neurotransmitter between P10 and P15.

**Figure 1:**
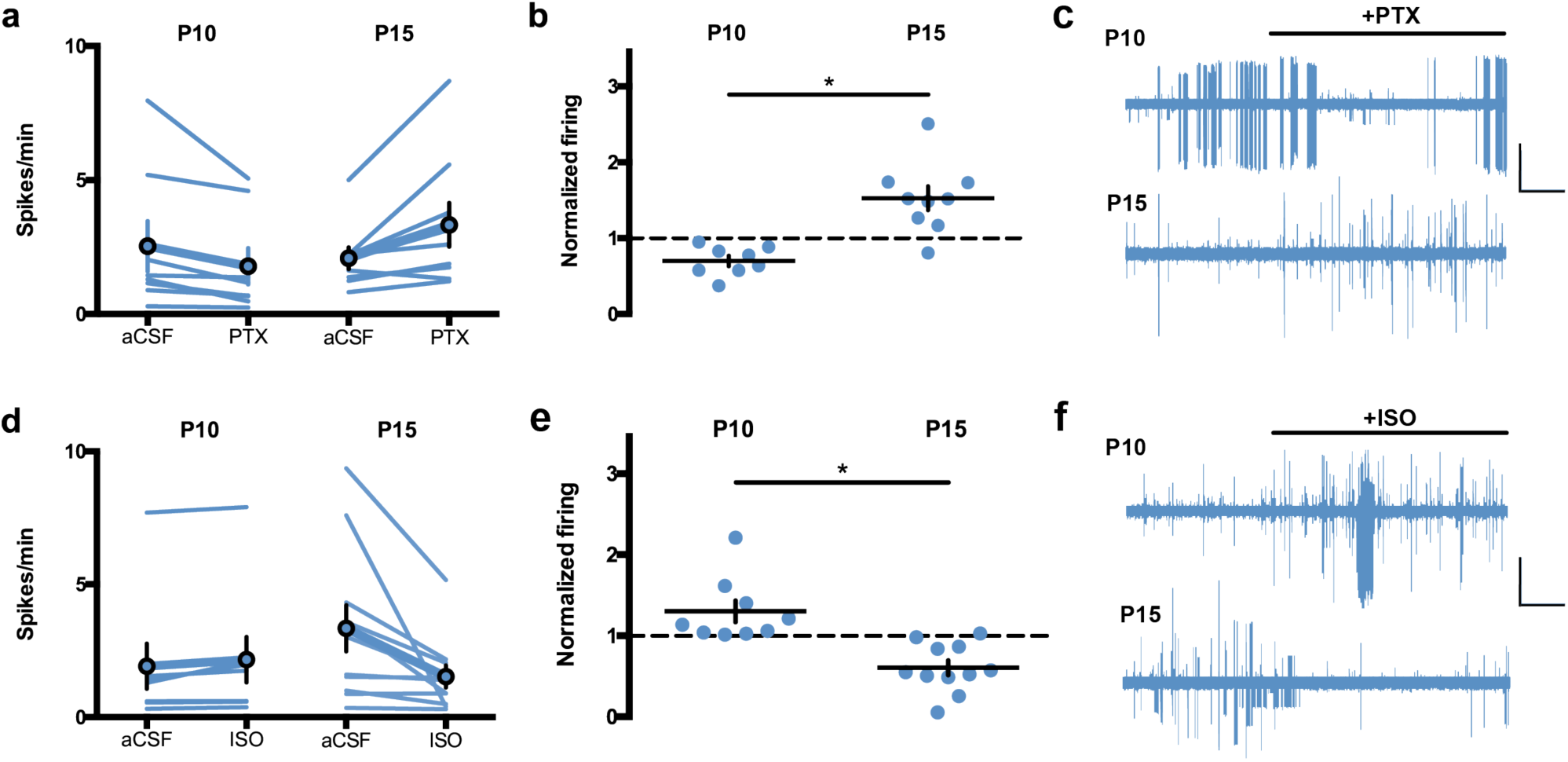
Developmental shift from excitation to inhibition by GABA-A receptors in rat medial prefrontal cortex slices. Action potentials were recorded in cell-attached (I=0) layer 5 pyramidal neurons in standard aCSF. After 10 min of baseline recording, picrotoxin (20M; GABA-A receptor antagonist, PTX) or isoguvacine (7μ M; GABA-A receptor agonist, ISO) was bath-applied. Spiking activity was calculated as an average of spikes per minute (10 min baseline) compared to the last 10 min of drug application. **a, b**: GABA-A receptor antagonism is inhibitory in immature P09-P10 PFC networks. PTX decreased spike frequency in slices obtained from P09-10 rats (N=8). In contrast, PTX increased spike frequency in slices obtained from P15-16 rats (N=9). **c**: Example traces (reference bars: 100pAx2min). **d, e**: GABA-A receptors agonism is inhibitory at P15-P16. ISO increased spike frequency in slices obtained from P09-10 rats (N=9). In contrast, ISO application decreased spike frequency in slices obtained from P15-16 rats (N=11). **f**: Example traces (reference bars: 100pAx2min). Error bars indicate SEM. Mann-Whitney test, *p<0.05.

### The GABA “switch” is delayed in pups perinatally exposed to WIN or THC

Next, to assess the impact of perinatal exposure through lactation to cannabinoids on this established trajectory of the PFC GABA transition, dams were injected with either WIN (0.5mg/kg s.c.) or THC (2mg/kg s.c.) from P01 to P10. Cell-attached recordings were then performed on layer 5 PFC neurons from their progeny at three time points (Figure 2). At P09-10, bath-application of PTX significantly reduced spike frequency in slices obtained from pups perinatally exposed to either WIN or THC (∼39% and 52% reduction, respectively; P<0.05 as compared to respective baselines) while application of ISO significantly increased spike frequency in both groups (∼53% and 49% increase, respectively; P<0.05 as compared to respective baselines). At P15-16, bath-application of PTX still attenuated spike frequency in slices obtained from WIN- or THC-exposed progeny (∼39% and 42% reduction, respectively; P<0.05 as compared to respective baselines). Similarly, ISO continued to increase spike frequency at this time point in both groups (∼40% and 53% increase, respectively; P<0.05 as compared to respective baselines). Thus, it can be concluded that GABA remains excitatory at P15-P16 in pups perinatally exposed to cannabinoids.

**Figure 2:**
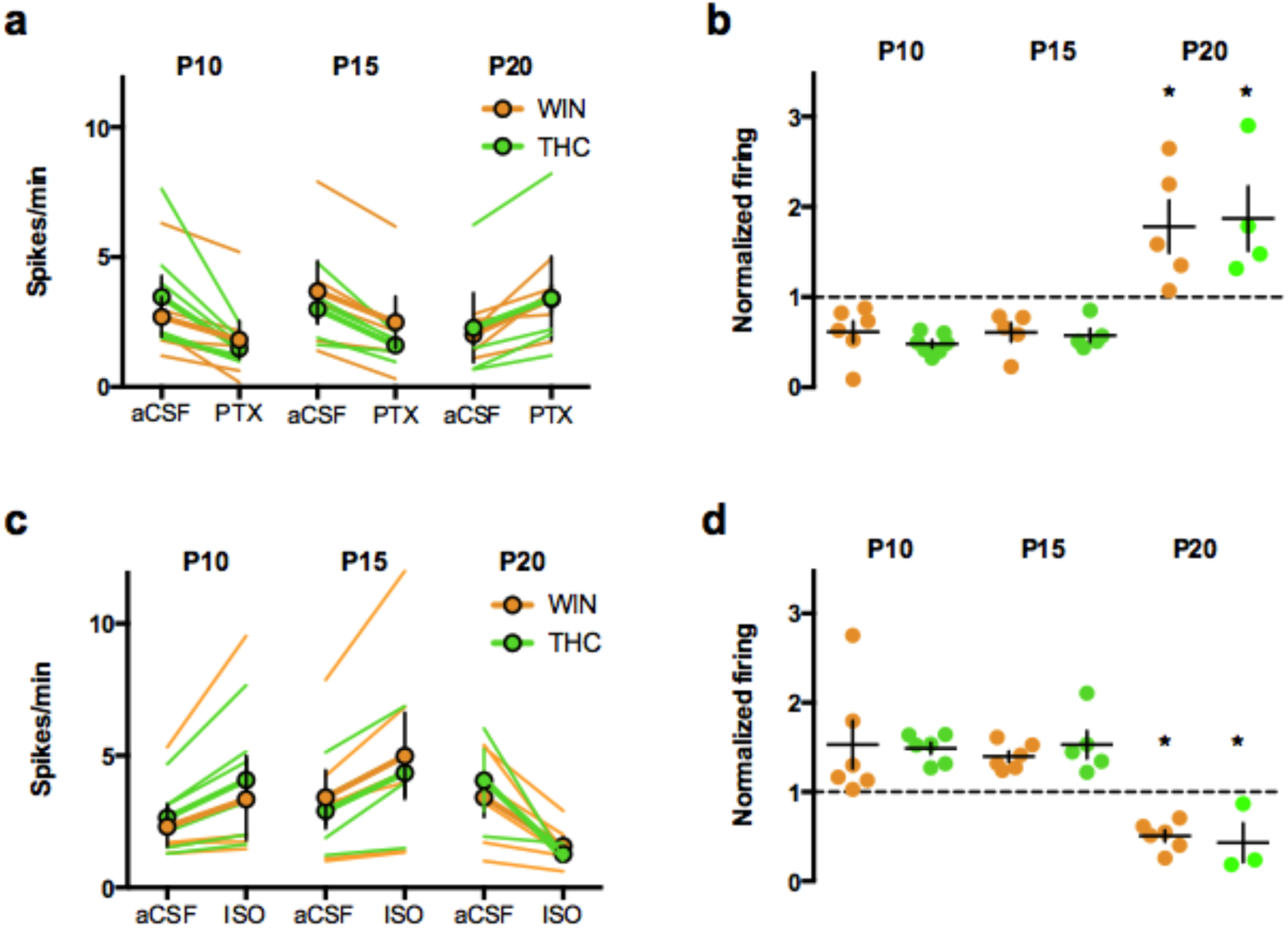
The excitatory/inhibitory switch is delayed in progeny of dams exposed to WIN or THC during lactation. Action potentials were recorded in cell-attached (I=0) layer 5 pyramidal neurons in standard aCSF. After 10 min of baseline recording, picrotoxin (20μ M; GABA-A receptor antagonist, PTX) or isoguvacine (7μ M; GABA-A receptor agonist, ISO) was bath-applied. Spiking activity was calculated as an average of spikes per minute (10 min baseline) compared to the last 10 min of drug application. **a, b**: GABA-A receptor antagonism is inhibitory at P09-10 and P15-16 but not P20-21 in animals perinatally exposed to WIN or THC. At P09-10, PTX decreased spike frequency in slices obtained from WIN (N=6) and THC (N=7) rats. Similarly, PTX bath-application decreased spike frequency in slices obtained from WIN (N=5) and THC (N=5) rats. However, at P20-21, PTX increased spike frequency in both WIN (N=5) and THC (N=4) rats. **c, d**: At P09-10, ISO increased spike frequency in WIN (N=6) and THC (N=6) rats. Similarly, at P15-16, ISO bath-application increased spike frequency in Perinatal WIN (N=6) and THC (N=5) rats. However, at P20-21, ISO decreased spike frequency in both WIN (N=6) and THC (N=3) rats. Two-way ANOVA followed by Sidak’s multiple comparisons test, F_5,26_ = 10.42, p<0.001. Error bars indicate SEM. *p<0.05 as compared to P09-10 or P15-16 of corresponding group.

Considering that the GABA switch is delayed in a number of identified disorders (Tang et al. 2016; He et al. 2014; Amin et al. 2017) as well as alterations to maternal health (Corradini et al. 2017) or behavior (Furukawa et al. 2017), we elected to perform recordings on slices obtained from progeny of the WIN- or THC-treated dams at P20-21 in order to ascertain whether the GABA switch had occurred at this age. Interestingly, PTX application at this time point elicited an increase in spike frequency in slices obtained from pups originating from either WIN- or THC-treated dams (∼78% and 87% increase, respectively; P<0.05 as compared to respective baselines). Additionally, bath-application of ISO elicited a significant decrease in spike frequency in slices obtained from these WIN- or THC-exposed pups (49% and 57% decrease, respectively; P<0.05 as compared to respective baselines). Together with the previous results, these findings indicate that in pups perinatally exposed to cannabinoids, GABA’s transition from an excitatory to inhibitory neurotransmitter is delayed, rather than entirely absent.

### The delayed GABA “switch” in perinatally cannabinoid-exposed pups is mediated by CB1R

In order to determine the mechanism underlying the effect of perinatal cannabinoid exposure on GABAergic development, dams were co-administered the CB1R antagonist AM251 along with WIN from P01-P10 (Figure 3). Cell-attached recordings of slices obtained from their progeny revealed a CB1R dependence in the previously identified WIN-induced retardation of the GABA switch. Specifically, at P09-10, bath-application of PTX induced a significant reduction in spike frequency (∼73%; P<0.05 as compared to baseline) while ISO significantly increased spike frequency by ∼38% (P<0.05 as compared to baseline). In line with our previous findings that GABA becomes an inhibitory neurotransmitter in the PFC by P15, slices obtained from the progeny of AM+WIN-treated dams at P15-16 exhibited increased spike frequency following bath-application of PTX (∼82%; P<0.05 as compared to baseline). Similarly, application of ISO decreased spike frequency at this time point by ∼50% (P<0.05 as compared to baseline). Together, these results reveal a CB1R-dependant mechanism for the perinatal WIN-treatment induced delay of GABA’s transition from an excitatory to inhibitory neurotransmitter.

**Figure 3:**
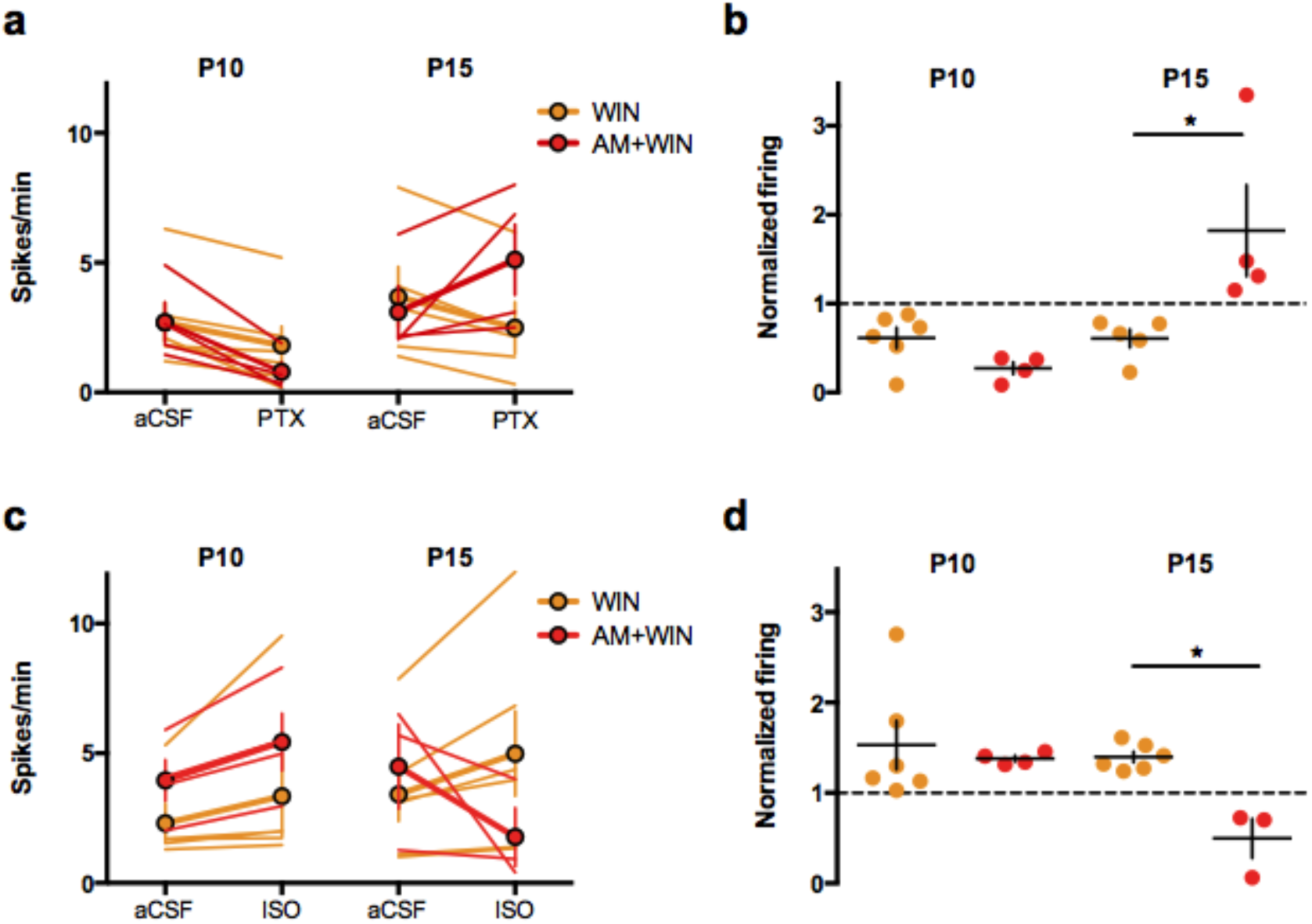
The delayed excitatory/inhibitory switch in progeny of dams exposed to WIN or THC is mediated by CB1R. Action potentials were recorded in cell-attached (I=0) layer 5 pyramidal neurons in standard aCSF. After 10 min of baseline recording, picrotoxin (20μ M; GABA-A receptor antagonist, PTX) or isoguvacine (7μ M; GABA-A receptor agonist, ISO) was bath-applied. Spiking activity was calculated as an average of spikes per minute (10 min baseline) compared to the last 10 min of drug application. AM251 co-administration prevents the delayed GABA shift induced by perinatal WIN treatment. **a, b**: GABA-A receptor antagonism is inhibitory at P09-10 and P15-16 in WIN progeny but only at P09-10 in AM+WIN. At P09-10, PTX decreased spike frequency in slices obtained from WIN (N=6) and AM+WIN (N=4) rats. At P15-16, PTX decreased spike frequency in WIN-but not AM+WIN-exposed progeny (N=5, 4 respectively). **c, d**: At P09-10, ISO application increased spike frequency in WIN (N=6) and AM+WIN (N=4) rats. However, at P15-16 ISO application increased spike frequency in slices obtained from WIN-exposed progeny (N=6) but decreased spiking in slices obtained from AM+WIN-exposed rats (N=3). Two-way ANOVA followed by Sidak’s multiple comparisons test, F_9,47_ = 10.46, p<0.0001. Error bars indicate SEM. *p<0.05.

### KCC2 upregulation is delayed in perinatally cannabinoid-exposed pups

The previously identified GABA “switch” is surmised to result from increased expression of KCC2 as identified in the hippocampus and elsewhere (Kaila et al. 2014; Valeeva, Valiullina, and Khazipov 2013). In addition to upregulation of KCC2, expression levels of NKCC1 also vary during the developmental GABA “switch.” We therefore elected to measure KCC2 and NKCC1 levels at our previously identified functionally relevant time-points for the GABA transition (Figure 4). First, we found that KCC2 levels significantly increase in the PFC of pups raised by vehicle-treated dams between P10 and P15. However, KCC2 levels remain unchanged between P10 and P15 in rats perinatally exposed to WIN. By P21, levels of KCC2 in WIN-exposed pups are significantly elevated to a level on par with P15 pups from sham-treated dams. Conversely, NKCC1 levels exhibit a decreasing trend in the PFC of rat progeny of sham-treated dams. As with KCC2, levels of NKCC1 remain unchanged in the PFC of WIN-exposed pups between these time points. Furthermore, the level of NKCC1 in the PFC of the WIN-exposed progeny remains unchanged at P21. Together, these data indicate that at P15, the lack of an apparent GABA transition from an excitatory to inhibitory role in the PFC of WIN-exposed pups may be attributed to a failure of KCC2 up regulation. Additionally, the delayed GABA “switch” which occurs by P21 in WIN-exposed progeny correlates with a significant increase in KCC2 levels.

**Figure 4:**
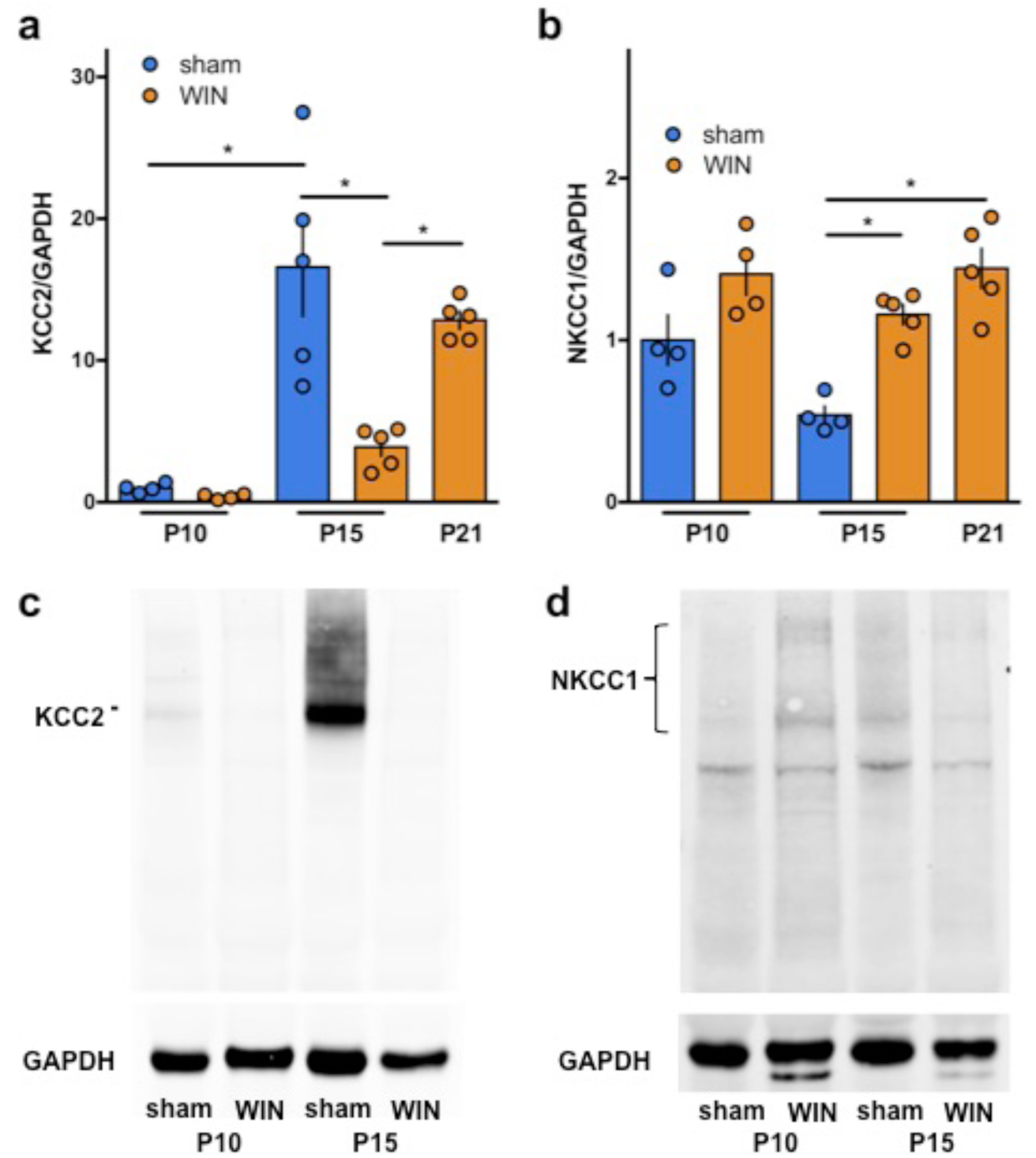
Perinatal WIN-exposure alters the developmental trajectory of KCC2 and NKCC1 expression in the PFC. Western-blot analysis of KCC2 and NKCC1 reveal altered expression levels between P10, P15 and P21 in progeny of dams exposed to WIN during lactation as compared to progeny of sham-treated dams. **a**: KCC2 levels are significantly increased between P10 and P15 in the PFC tissue collected from pups of sham-treated dams (P10 N=4, P15 N=5). However, no change in KCC2 levels was detected in PFC tissue collected from pups of WIN-treated dams (P10 N=4, P15 N=5). At P21, a significant increase in KCC2 was observed in PFC tissue from WIN-treated pups (N=5). One-way ANOVA followed by Sidak’s multiple comparisons test, F_4,18_ = 5.813, p<0.0001. **b**: At P10, no difference in NKCC1 levels was detected in PFC tissue collected from pups of sham- or WIN-treated dams was detected. However, at both P15 and P21, PFC tissue from pups of WIN-treated dams (P15 N=5, P21 N=5) revealed significantly higher levels of NKCC1 as compared to pups of sham-treated dams (P15 N=4). One-way ANOVA followed by Sidak’s multiple comparisons test, F_4,17_ = 0.9624, p=0.0002. Error bars indicate SEM.*p<0.05. **c,d**: Representative Western-blots of KCC2/GAPDH and NKCC1/GAPDH, corresponding to **a,b** respectively. Representative blots of KCC2/GAPDH and NKCC1/GAPDH for P10/P15/P21 of WIN-exposed progeny may be found in supplemental Figure 1.

**Figure 5:**
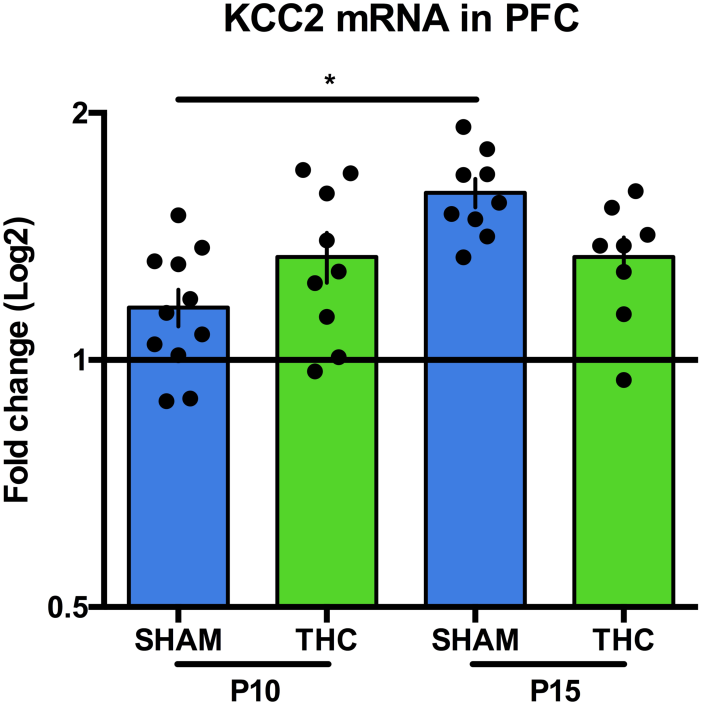
Perinatal THC exposure alters the developmental trajectory of KCC2 mRNA. qPCR analysis of KCC2 mRNA reveal altered expression levels between P10 and P15 in progeny of dams exposed to THC during lactation as compared to progeny of sham-treated dams. **a**: Levels of KCC2 mRNA are significantly increased between P10 and P15 in PFC tissue collected from pups of sham-treated dams (P10 N=11, P15 N=9). However, no change in KCC2 mRNA levels was detected in PFC tissue collected from pups of THC-treated dams (P10 N=9, P15 N=8). One-way ANOVA followed by Sidak’s multiple comparisons test, F_3,33_ = 6.495, p=0.0014. Error bars indicate SEM. *p<0.05

### KCC2 mRNA transcriptional upregulation between P10 and P15 in perinatally cannabinoid-exposed pups

In order to determine whether the delayed upregulation of KCC2 observed in pups perinatally exposed to exogenous cannabinoids occurs as a result of alterations in transcription or translation, we elected to perform qPCR on brains extracted from THC-exposed pups at the previously identified pertinent time points. First, we found that in the PFC of pups from sham-treated dams, mRNA for KCC2 is significantly increased between P10 and P15. However, mRNA for KCC2 remains unchanged between these two time points in the PFC of pups from THC-treated dams. These results indicate that exogenous cannabinoid exposure during lactation induces an alteration in KCC2 expression at least in part at the transcriptional level.

### The delayed GABA “switch” in perinatally cannabinoid-exposed pups is prevented by bumetanide treatment

Inhibition of NKCC1 has been studied as a potential pharmacotherapeutic means of correcting problems in GABAergic development in such conditions as neonatal seizures (Löscher et al. 2013) and maternal separation-induced stress (Hu et al. 2017), amongst others in which GABA maintains an excitatory role. Thus, to confirm the role of the KCC2/NKCC1 imbalance in the effects of perinatal WIN exposure, we treated pups with the NKCC1 inhibitor bumetanide from P01 to P15 during concomitant exposure to WIN via lactation. As before, cell-attached recordings were performed on slices obtained from these pups at P10 and P15 to determine the excitatory/inhibitory properties of GABA at these time points (Figure 6). At P10, bath-application of PTX decreased spike frequency indicating that GABA exhibited excitatory properties (∼58%; P<0.05 as compared to baseline). In accordance with this finding, bath-application of ISO increased the firing rate of cells recording at this time point by ∼78% (P<0.05 as compared to baseline). On the contrary, bath-application of PTX on slices obtained from P15 bumetanide-treated pups increased firing frequency (∼18%; P<0.05 as compared to baseline) while ISO induced a decrease in the firing rate (∼41% P<0.05 as compared to baseline). Therefore, we can conclude that an imbalance of KCC2/NKCC1 plays a crucial role in the delayed GABA “switch” exhibited by pups perinatally exposed to WIN.

**Figure 6:**
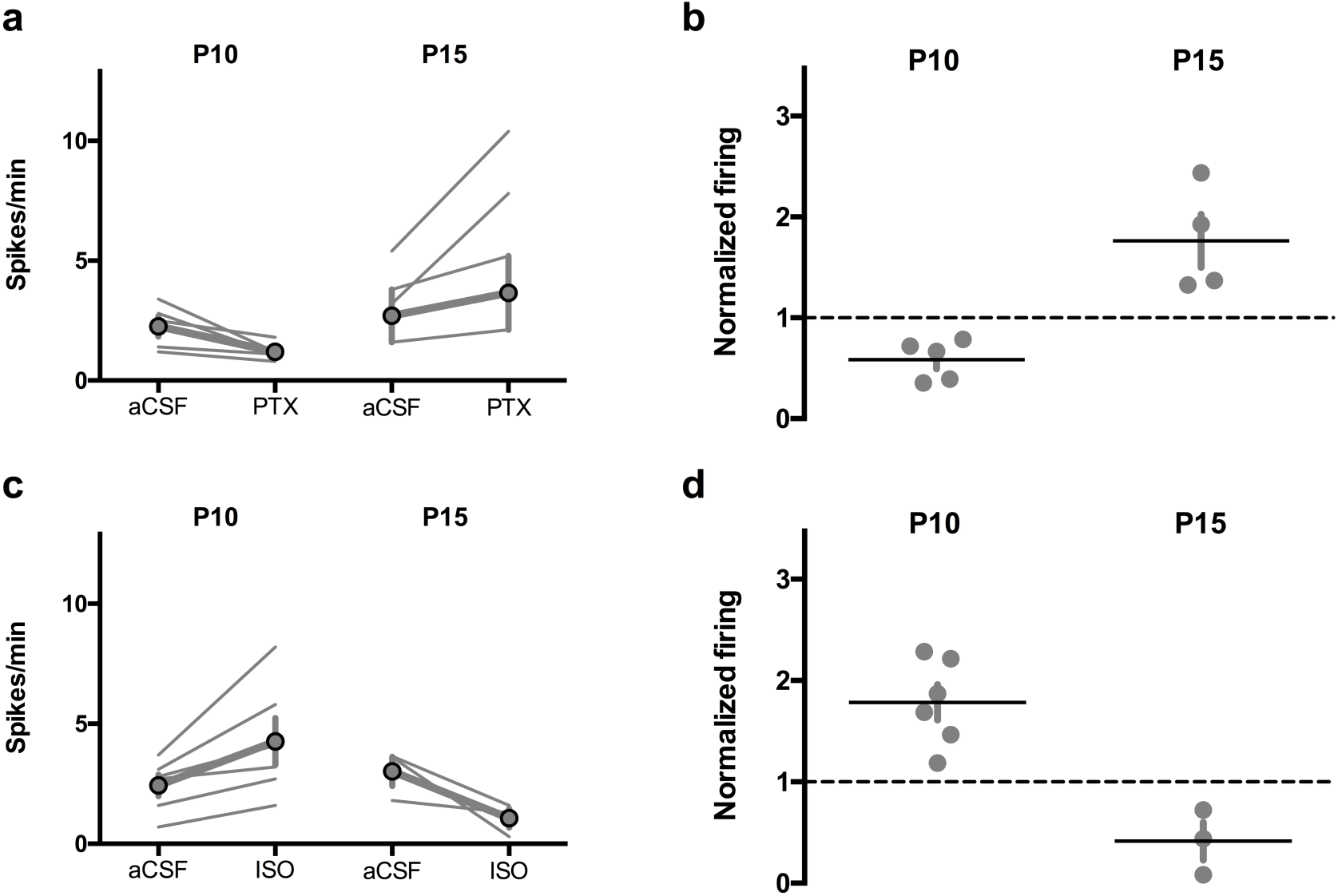
The delayed excitatory/inhibitory switch in progeny of dams exposed to WIN is prevented by perinatal (P1-P15) bumetanide treatment. Action potentials were recorded in cell-attached (I=0) layer 5 pyramidal neurons in standard aCSF. After 10 min of baseline recording, picrotoxin (20μ M; GABA-A receptor antagonist, PTX) or isoguvacine (7μ M; GABA-A receptor agonist, ISO) was bath-applied. Spiking activity was calculated as an average of spikes per minute (10 min baseline) compared to the last 10 min of drug application. **a, b**: GABA-A receptor antagonism is inhibitory at P09-10 in WIN+bumetanide, but excitatory at P15-16. At P09-10, PTX decreased spike frequency in slices obtained from WIN+bumetanide rats (N=5). Conversely, at P15-16 PTX decreased spike frequency in WIN+bumetanide-exposed progeny (N=4). Two-tailed T-test with Welch’s correction, P=0.0156. **c, d**: At P09-10, ISO application increased spike frequency in WIN+bumetanide rats (N=6). However, at P15-16 ISO application increased spike frequency in slices obtained from WIN+bumetanide exposed progeny (N=3). Two-tailed T-test with Welch’s correction, P=0.0023.

## Discussion

The developmental consequences of perinatal cannabinoid exposure remain woefully under researched, despite the increasing availability of cannabis and its reported use by individuals both during and following pregnancy. Here, we sought to identify the potential implications of cannabinoid exposure in early development by treating lactating dams with either a synthetic cannabinoid (WIN) or the main psychoactive ingredient of cannabis-based preparations (THC), followed by cell-attached electrophysiological assessment of GABAergic function in the PFC in addition to measuring expression levels of relevant proteins in this region.

First, our results revealed that GABA does indeed exhibit excitatory properties in layer 5 of the PFC in early development, before transitioning to an inhibitory role. This « switch » occurs between P10 and P15, ascertained by cell-attached recordings of pyramidal neurons in layer 5 of the PFC. These results are in line with the timing of this transition identified in previous studies in other regions of the developing rat brain, including the hippocampus (Rivera et al. 1999; Tyzio et al. 2006) and cerebellum (Ben-Ari 2008) as well as in layer 2/3 cells of the neocortex (Luhmann and Prince 1991). Additionally, Western blot and qPCR analyses demonstrated that this switch is driven by an increase in KCC2 mRNA and protein, in parallel with findings in other brain regions (Ben-Ari et al. 2007).

The present results showed that maternal exposure to cannabinoids retards GABAergic development in the PFC. Pups originating from dams treated with either WIN or THC during the first 10 days of postnatal development exhibit a significant and CB1-dependent delay in the PFC GABA “switch.” This lack of a functional transition from an excitatory to inhibitory role for GABA was found to be associated with a suppressed trajectory of KCC2 protein and mRNA levels during this developmental period. In addition to previous research regarding the importance of this period in developing chloride homeostasis via regulation of the transport proteins KCC2 and NKCC1, this period has been recently identified as a crucial time-point in the development of GABAergic synapse innervation of the PFC (Virtanen et al. 2018), further underscoring the relevance of this vulnerable developmental trajectory with regards to GABA function. Interestingly, the delayed “switch” was prevented by direct administration to the developing pups of the NKCC1 antagonist bumetanide, which decreases intracellular chloride (**ref**).

Currently, the long-term consequences of this delay are unknown, however retarded GABA development has been associated with significant disorders of neuronal function that contribute to developmental aberrations such as Fragile X syndrome (He et al. 2014), 22q11.2 deletion syndrome (Amin et al. 2017), early life epilepsies (Briggs et al 2011) and autism (Cellot and Cherubini 2014; Sgadò et al. 2011; Tyzio et al. 2006). In addition, there is precedence for alterations in GABAergic development resulting from a maternal insult such as immune activation (Corradini et al. 2017). However, we present here the first data suggesting that postnatal development of GABAergic signaling may be altered through drug-exposure during lactation.

GABAergic development has significant consequences reaching beyond GABAergic synapses, including the regulation of synaptic integration of newborn neurons and the titration of glutamatergic signaling (Ge et al. 2006). Additionally, such developmental phenomena as neuronal proliferation, migration and synaptogenesis are similarly governed by developing GABAergic transmission (Ben-Ari 2002). Previous research indicates that alterations in the time-course of GABAergic development in cortical regions impacts excitatory glutamatergic transmission as well, which consequently exhibits as sensorimotor gating deficits associated with schizophrenia-like behavioral outcomes (Wang and Kriegstein 2011). The apparent retardation of the GABA development caused by perinatal cannabinoid exposure therefore likely impacts a wide array of neuronal functions in the PFC and elsewhere, whose impact in later stages of development remain to be investigated.

Together, our results indicate that the developmental trajectory of GABA in PFC neurons is significantly altered by exposure to cannabinoids through lactation during the early perinatal period. This aberrant development exhibits as a delayed GABA “switch” caused by a lack of KCC2 upregulation due to altered mRNA levels. Furthermore, the prevention of the delayed GABA “switch” by direct treatment of bumetanide in the developing pups confirms the mechanistic role of a KCC2/NKCC1 imbalance in this altered trajectory. However, this does not suggest a viable strategy for treatment of a developmental retardation that may be caused by perinatal cannabinoid exposure, as bumetanide treatment in the developing infant is associated with significant ototoxicity during this time period (Ben-Ari et al. 2016; Delpire et al. 1999). Further analyses of both electrophysiological function and its molecular underpinnings, as well as behavioral consequences associated with this aberrant development may reveal long-term consequences of these early postnatal developmental alterations.

## Methods

### Animals

Animals were treated in compliance with the European Communities Council Directive (86/609/EEC) and the United States National Institutes of Health Guide for the care and use of laboratory animals. All rats were group-housed with 12 h light/dark cycles and ad libitum access to food and water. All behavioral and synaptic plasticity experiments were performed on male and female RjHan:wi-Wistar rats between P09 and P21. Male and female electrophysiological and biochemical results exhibited no difference; thus the data were pooled.

### Drug treatments

Sub-cutaneous injections were performed daily from postnatal day (PND) 01 to PND10 with the synthetic cannabinoid WIN55,212-2 (WIN; 0.5 mg/kg) alone or in combination with AM-251 (0.5mg/kg), or with the phytocannabinoid Delta9-Tetrahydrocannabinol (THC; 2 mg/kg). WIN55,212-2 was suspended in 5% DMSO, 5% cremophor and saline, at a volume of 1 ml/kg. Control dams (Sham group) received a similar volume injection of vehicle solution. THC was suspended in 5% Ethanol, 5% cremophor and saline, and injected s.c. at a volume of 1 ml/kg. Control dams (SHAM group) received a similar volume injection of vehicle solution. Newborn litters found up to 05:00 p.m. were considered to be born on that day. Bumetanide treatments were performed twice daily (09:00 a.m. and 05:00 p.m.) from P1 to P15. Pups were injected with 0.2mg/kg bumetanide in 0.1% DMSO, 99.9% saline at a volume of 10µl/g.

### Electrophysiology

Coronal slices containing the prelimbic area of the medial prefrontal cortex (PrPFC) were prepared as previously described (Lafourcade et al., 2007). Briefly, mice were anesthetized with isoflurane and 300µm-thick coronal slices were prepared in a sucrose-based solution (in mM: 87 NaCl, 75 sucrose, 25 glucose, 2.5 KCl, 4 MgCl2, 0.5 CaCl2, 23 NaHCO3 and 1.25 NaH2PO4) at 4°C using an Integraslice vibratome (Campden Instruments). Slices were stored for one hour at 32°C in artificial cerebrospinal fluid (ACSF; in mM: 130 NaCl, 2.5 KCl, 2.4 MgCl2, 1.2 CaCl2, 23 NaHCO3, 1.2 NaH2PO4 and 11 glucose), equilibrated with 95%O2/5% CO2. Slices were then stored at room temperature until recording. All experiments were conducted at 30–32°C in ACSF. Cell-attached patch-clamp recordings were made in PrPFC layer5/6, collected using an Axopatch-1D amplifier (Axon Instruments) and acquired with Clampex 10.2 acquisition Software via a Digidata 1440A (Axon Instruments). Pyramidal neurons in PrPFC layer 5/6 were visualized using an infrared illuminated upright microscope (Olympus BX51WI). Slices were superfused at 2ml/min with aCSF.

### Spontaneous Spiking Activity

Spontaneous spiking activity was recorded in cell-attached configuration with a patch pipette filled with ACSF. A >500MOhm seal was obtained in current clamp configuration before recording spike-activity in I D0 mode. Data were filtered at 2kHz and digitizedat 10kHz. Spontaneous spike activity was analyzed in Clampfit 10.5 (MolecularDevices) threshold detection with a trigger threshold of >2x SD of baseline noise. Mean spike activity was calculated as an average of spikes per minute over a 10-minute baseline period. For drug-effects,mean spike activity was calculated as an average of spikes per minute over a 10-minute period following at least 5 minutes of bath perfusion.

### Western-blots

Animals were anesthetized with isoflurane and brains were rapidly removed and dissected before being placed in liquid nitrogen and stored at -80°C until further processing. For Western blot processing, after dissecting the mPFC from 1 mm coronal sections, samples were homogenized in RIPA buffer (NaCl 150mM; triton X-100 1% deoxycholate 0.5%; SDS 0.1% Tris HCl 50mM pH 8.0) and centrifuged at 10,000xg for 10 minutes at 4°C. The supernatant was then mixed with 4X sample buffer and incubated for 10 minutes at 65°C and run on a 4-12% NuPage gel (Thermo Fisher Scientific, Florence, KY). Following protein transfer, blots were stained with Ponceau S to compare protein loading and then blocked in LiCor Blocking Buffer (LiCor Bioscience, Lincoln, NE) for 60 minutes at room temperature. They were then incubated with either rabbit anti-KCC2 (1:2,000; NeuroMab, UC Davis, CA, Catalog # 75-050) or rabbit anti-NKCC1 (1: 1,000; Cell Signaling Technology, Danvers, MA, Catalog number 14581) diluted in a mixture of LiCor Blocking Buffer and 1XPBS (1:1). At the same time, blots were incubated with mouse anti-GAPDH (1:1,000; Biolegend, San Diego, CA, Catalog number 649201, clone number FF26A/F9). Blots were incubated with these primary antibodies overnight at 4°C. The next day, blots were washed 4 × 15 minutes at room temperature in TBST (20 mM Tris, pH 7.5, 150 mM NaCl, 0.05% Tween 20) and then incubated in LiCor Blocking Buffer and 1XPBS (1:1) containing donkey anti-rabbit IR680 and donkey anti-mouse IR800 antibodies (1:10,000; LiCor, NE, Catalog#s 926-68073, 926-32212, rabbit and mouse respectively) for 1 hour at room temperature. Finally, blots were washed as above and scanned on a LiCor Odyssey near-IR imager. Band densities were collected using FIJI software and those corresponding to KCC2 and NKCC1 were normalized to GAPDH density.

### qPCR

Medial prefrontal cortex was harvested from frozen rat brains via a coronal cutting block (Braintree Scientific, Inc., Braintree, MA, Cat.# BS-SS 605C). The block and glass plate used for dissection had been pre-chilled to <-20°C and this temperature was maintained throughout the process using pelleted dry ice. Harvested mPFCs were stored at -80°C until use. Total RNA was extracted using the RNeasy Plus Micro Kit (Qiagen, Hilden, Germany, Cat.# 74034) and Reverse transcription was performed using the RevertAid Kit (Thermo Fisher Scientific, Waltham, MA Cat.# K1621) as per manufacturer’s instructions. Taqman primers and probes were obtained through Applied Biosystems. Sequences used are as follows:

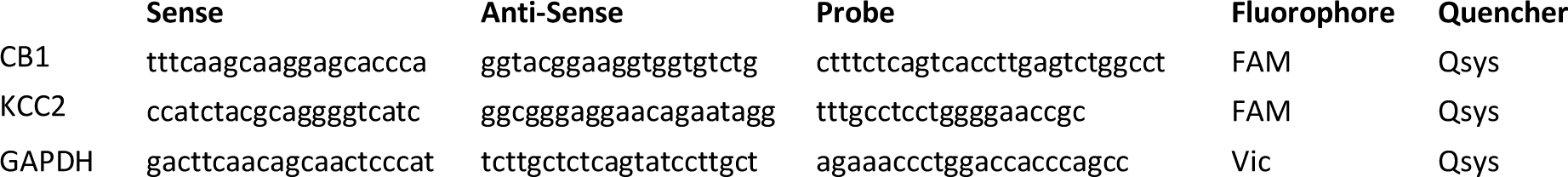

Gene expression data was generated on a QuantStudio7 thermal cycler using the TaqMan Gene Expression Master Mix from Applied Biosystems (Foster City, CA Cat.# 4369016). Duplicates were run for each sample, and relative gene expression was determined using the ⍰⍰Ct method. Excel (Microsoft, Redmond, WA) and Prism 7 (Graphpad, La Jolla, CA) software was used for data analysis.

### Statistics

Analyses were conducted using Microsoft Excel 2016 (Redmond, WA) and GraphPad Prism 7 (La Jolla, CA), with the level of significance set at 0.05. Significance was assessed by one-way analysis of variance (ANOVA), or by Student’s t-test if two samples were compared. If a significant difference in ANOVA was detected, Sidak’s multiple comparisons post hoc analyses were conducted.

## Acknowledgments

The authors are grateful to Dr. Christophe Pelegrino for help and advice with the western blot experiments, Drs. Pascale Chavis, Camille and Jean-Luc Gaiarsa for helpful discussions and to the National Institute of Mental Health’s Chemical Synthesis and Drug Supply Program (Rockville, MD, USA) for providing CNQX and SR141716A.

